# Epigenetics of post-operative delirium: A genome-wide DNA methylation study of neurosurgery patients

**DOI:** 10.1101/2022.10.04.510706

**Authors:** Takehiko Yamanashi, Kaitlyn J. Crutchley, Nadia E. Wahba, Takaaki Nagao, Pedro S. Marra, Cade C. Akers, Eleanor J. Sullivan, Masaaki Iwata, Mathew A Howard, Hyunkeun R. Cho, Hiroto Kawasaki, Christopher G. Hughes, Pratik P. Pandharipande, Marco M. Hefti, Gen Shinozaki

**Author notes:** Takehiko Yamanashi, Kaitlyn J. Crutchley, and Nadia E. Wahba are co-first authors. **Corresponding Author**, Gen Shinozaki, Stanford University School of Medicine, Department of Psychiatry and Behavioral Sciences, 3165 Porter Drive Room 2175, Palo Alto, CA, 94304, USA. **Previous presentation:** none. **Disclosures:** Corresponding author, Gen Shinozaki has pending patents “Epigenetic Biomarker of Delirium Risk” in the PCT Application No. PCT/US19/51276, and in U.S. Provisional Patent No. 62/731,599. All other authors have declared that no conflict of interest exists.

## Abstract

**Aims:** There is no previous study demonstrating the differences of genome-wide DNA methylation (DNAm) profiles between patients with and without postoperative delirium (POD). We aimed to discover epigenetic (DNAm) markers that are associated with POD in blood obtained from patients before and after neurosurgery.

**Methods:** Pre- and post-surgical blood DNA samples from 37 patients, including 10 POD cases, were analyzed using the Illumina EPIC array genome-wide platform. We examined DNAm differences in blood from patients with and without POD. Enrichment analysis with Gene Ontology and Kyoto Encyclopedia of Genes and Genomes terms were also conducted.

**Results:** When POD cases were tested for DNAm change before and after surgery, enrichment analyses showed many relevant signals with statistical significance in immune response related-pathways and inflammatory cytokine related-pathways such as “cellular response to cytokine stimulus”, “regulation of immune system process”, “regulation of cell activation”, and “regulation of cytokine production”. Furthermore, after excluding the potential effect of common factors related to surgery and anesthesia between POD cases and non-POD controls, the enrichment analyses showed significant signals such as “immune response” and “T cell activation”, which are same pathways previously identified from an independent non-surgical inpatient cohort. Conclusions: Our first genome-wide DNAm investigation of POD showed promising signals related to immune response, inflammatory response and other relevant signals considered to be associated with delirium pathophysiology. Our data supports the hypothesis that epigenetics are playing an important role in pathophysiological mechanism of delirium and suggest the potential usefulness of epigenetics based biomarker of POD.

## Introduction

Delirium, especially after surgery, is dangerous and common among elderly patients, yet it is underdiagnosed and undertreated (1–3). It is also associated with long-term cognitive decline (4) and high mortality (5). As such, it is becoming increasingly important to predict which surgical patients are at risk for delirium and to better understand the pathophysiological mechanisms of this condition. One way this can be achieved is with the use of biomarkers (6–9). Previous studies have revealed that elderly patients with delirium have increased serum levels of inflammatory markers and cytokines (6–9). These data suggest that changes in the molecular mechanism for increased expression of cytokine genes can be detected and can serve as biomarkers to identify individuals with risk for delirium (10). It is noteworthy that a well-known risk factor for delirium is aging, although the molecular process of how aging may enhance cytokine release and make individuals more vulnerable to delirium is not well understood. Understanding this mechanism is crucial to better understand pathophysiological mechanisms of delirium and to improve clinical practice, including prevention and therapeutic intervention.

Of note, DNA methylation (DNAm) is well known to be tightly regulated with aging (11–13). In addition, DNAm is important epigenetics machinery for controlling expression of genes across the genome (14). These facts suggest that DNAm can be a promising target for investigating the molecular mechanism of delirium and has the potential to serve as a biomarker for the condition (10).

The literature related to the role of epigenetics in delirium is sparse (10). Our group has investigated the differences of epigenetics (especially DNAm) status between delirium cases versus controls (15–17), and we previously showed that the DNA methylation in the pro-inflammatory cytokine gene, *TNF-alpha*, universally decreases with age among patients with delirium, but not in age- and sex-matched controls (17). In addition, we found that DNAm of the neurotrophic factor gene *BDNF* increases with age among delirium cases more significantly than controls (16). Also, enrichment analysis based on genome-wide data showed top signals indicating the role of immune response, inflammatory response, and cholinergic synaptic function in delirium (15).

Delirium cases from our prior work were from a heterogenous cohort with multiple different medical and surgical conditions leading to their hospitalization. Thus, it was not feasible to control for the etiology of their delirium. Also, biological samples prior to their delirium onset were not available, and we were not able to determine if the epigenetic signals we found among delirium subjects were due to the status of delirium itself, due to the baseline cause of delirium such as infection or surgery, or due to their baseline epigenetic marks prior to the development of delirium.

To overcome the limitations of our previous reports, in the present study, we aimed to discover epigenetic (DNAm) markers that are associated with postoperative delirium (POD) in blood obtained from patients before and after neurosurgery for brain resection as a treatment for medication-refractory epilepsy patients. Our hypothesis is that DNAm levels change across the genome specific to surgical insult and/or delirium occurs in blood, and that these DNAm signals confer phenotypic changes that exacerbate reactions to exogenous insult leading to delirium.

We specifically hypothesized that genome-wide DNAm signals differ between patients who experience POD after neurosurgery relative to those who do not. We tested this hypothesis by analyzing whole blood from a set of 37 neurosurgery patients aged 5 to 64 years. We aimed to identify 1) DNAm signals that are predictive of POD by comparing pre-surgical samples from patients with POD to those from those without POD. We also tested this set 2) for cross-sectional analysis using blood samples obtained post-surgery by comparing patients who experienced POD and patients who did not; and 3) for prospective analysis using post-surgical vs. pre-surgical samples from the POD group versus the non-POD group. By comparing these two pre/post datasets, we aimed to capture postoperative DNAm changes that are unique to POD cases excluding the influence from surgical procedure itself, including exposure to anesthesia.

## Methods

### Subject recruitments

Details of study subjects and the enrollment process have been described previously (18, 19). Briefly, we recruited subjects who were scheduled for brain resection surgery due to their medication-refractory epilepsy at the University of Iowa Hospitals and Clinics between April 2015 and July 2019. Written informed consent was obtained. If the subject was a minor, consent was obtained from their parents. This study was approved by the University of Iowa’s Human Subjects Research Institutional Review Board.

### Clinical categorization and data collection

Details of clinical data collection methods have been described in our previous publications (18, 19). Briefly, we obtained medical and surgical history and demographic information from electronical medical records and patient interviews. Detailed chart review was employed to identify evidence of delirium during the post-operative period (through 7 days after brain resection surgery, or until discharge, whichever came first) (20). POD-positive cases were identified based on if fluctuations in mental status including varied states of awareness and orientation via physical, neurological, and mental status exams were recorded. Additional data contributing to determination of categorization included Confusion Assessment Method—Intensive Care Unit (CAM-ICU) scores when available. Each case in question for grouping was reviewed by a board-certified consultation-liaison psychiatrist (G.S.) for final decision of POD categorization.

### Blood sample collection and processing

Whole blood was obtained using EDTA tubes before and after surgery. Blood samples were collected at the beginning of surgery as a “pre” sample and at the end of surgery as a “post” sample in the operating room (OR). In total, pre and post sets of blood samples were available for 37 subjects. Samples were stored at −80 °C until downstream DNA extraction and DNAm analysis.

### DNA methylation analysis

Genomic DNA was extracted from whole blood tissues with the MasterPure DNA extraction kit (MCD85201, Epicentre, Madison, WI, USA) following the recommended protocol. DNA quality was assessed with NanoDrop spectrometry and quantified with the Qubit dsDNA Broad Range Assay Kit (Q32850, ThermoFisher Scientific, Waltham, MA, USA). For each sample, 500□ng of DNA was bisulfite-converted with the EZ DNA Methylation Kit (D5002, Zymo Research, Irvine, CA, USA). The Infinium HumanMethylationEPIC BeadChip Kit (WG-317-1002, Illumina, San Diego, CA, USA) was used to analyze genome - wide DNAm. The arrays were scanned with the Illumina iScan platform.

The R packages ChAMP and Minfi were used to process the raw methylation data. During the loading of data, probes were filtered out if they (i) had a detection p-value >0.01, (ii) had <3 beads in at least 5% of samples per probe, (iii) were non-CpG, SNP related, or multi-hit probes, (iv) were located on chromosome X or Y. Samples were normalized with beta-mixture quantile dilation before performing differential methylation analyses.

### Statistical analysis

R was used for all statistical analysis (21). The chi-square test with Yates’ continuity correction was used to calculate the categorical data. Estimated cell proportions for CD8 T cells, CD4 T cells, natural killer cells, B cells, and monocytes were calculated by the DNAm Age Calculator available online (22, 23) through the method reported previously (24). DNA methylation differences at each CpG site were assessed by RnBeads using the limma method (25, 26). We tested DNAm differences between POD cases versus non-POD controls in 1) pre-surgery samples and 2) post-surgery samples. We also assessed DNAm changes after surgery by 3) the pre-post sample comparison [3a) total cohort, 3b) POD cases only, and 3c) non-POD controls only] (**Supplementary Figure 1**). Covariates included in the analysis were age, sex, and cell type proportions. Genome-wide significance was set at a p-value of less than 5.0E-08.

Gene Ontology (GO) and Kyoto Encyclopedia of Genes and Genomes (KEGG) terms were also evaluated for enrichment analysis using the gometh analysis with the R package missMethyl (27) by adjusting for the variable number of CpG sites tested in each gene. For each comparison extracted with limma, we included the top 1,500 CpG sites with p < 0.05 for GO and KEGG to capture potentially relevant pathways. As a part of the pre-post sample comparison described above, we also analyzed a set of differentially methylated CpG sites unique to the pre-post POD case group. This subset of CpG sites was determined by the following process: i) POD case and non-POD control groups were filtered for CpG sites with p < 0.05, ii) CpG sites in common between the two subsets of significant CpG sites were determined, iii) the set of common CpG sites was removed from the set of significant POD case CpG sites. iv) From this new set of CpG sites unique to POD case, we included the top 1,500 CpG sites to detect perturbations in GO and KEGG pathways.

## Results

### Participant demographics

A total of 37 patients who were scheduled for brain resection neurosurgery and enrolled in this study had available blood samples for this analysis. Their average patient age was 32.8 years (SD = 15.3), 35.1% were female, and 97.3% were non-Hispanic white per self-report. Among them, 10 patients developed POD, and 27 did not. Average age for POD group was 41.1 years (SD = 15.7); 5 of 10 (50.0%) were female. Average age for non-POD group was 29.8 years (SD = 14.3); 8 of 27 (29.6%) were female. There were significant differences in age between patients with and without POD (**Table 1**).

**Table 1.**
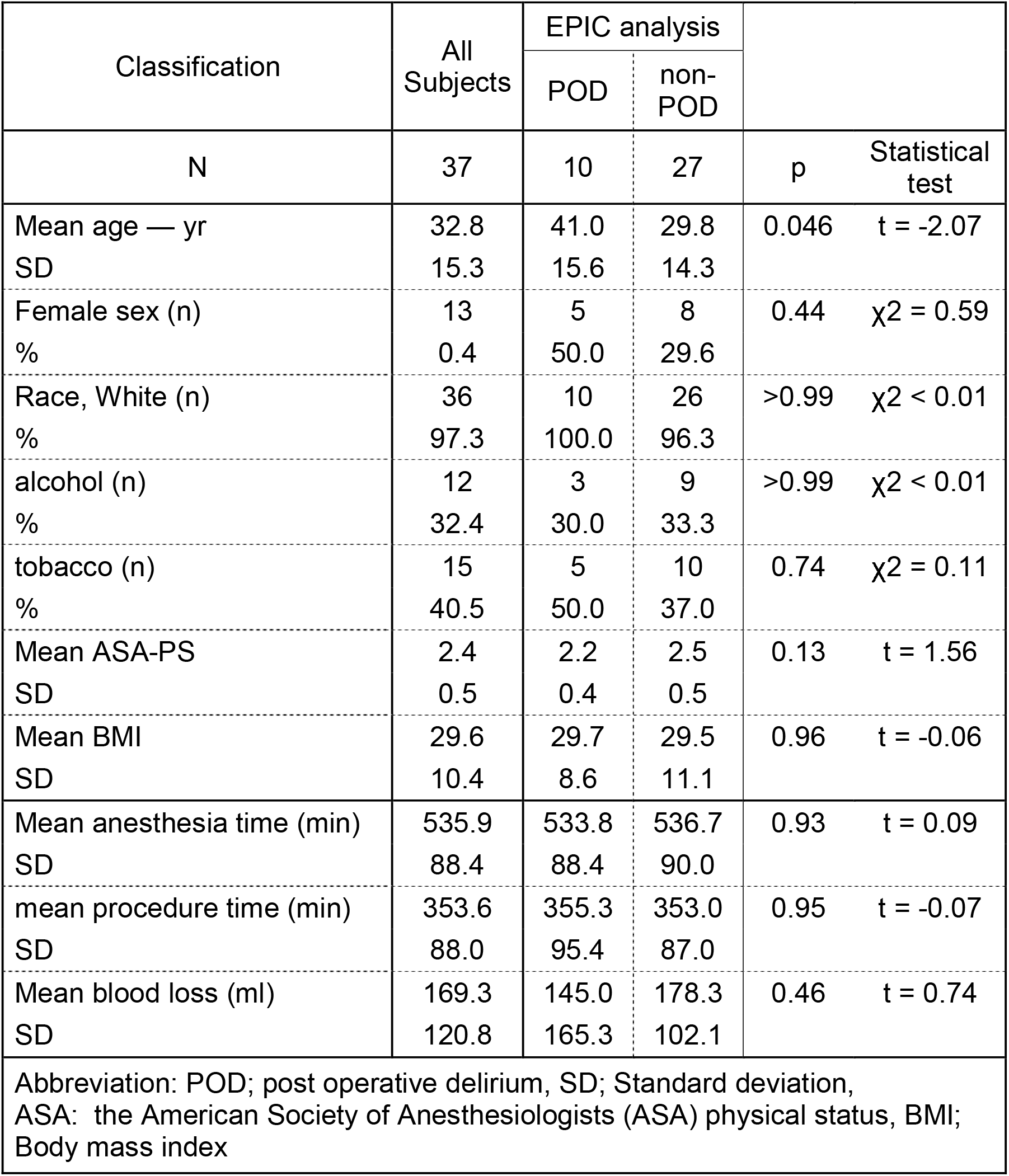
Patient Characteristics.

### POD case versus non-POD control Epigenome-wide association studies (EWAS) top hits

When pre-surgery samples were compared between POD and non-POD patients, no genome-wide significant signals were found (**Supplementary table 1**).

When blood samples from post-surgery were compared between POD cases and non-POD controls, no CpG signals were identified at the genome-wide significance level (**Supplementary table 2**).

When pre- and post-surgery samples were compared, no genome-wide significant signals were discovered with the total cohort, POD cohort only, or non-POD cohort only (**Supplementary table 3**).

### Enrichment analysis with GO and KEGG

1) Pre-surgery sample comparison between POD cases (n = 10) vs non-POD controls (n = 27): There were no significant pathways identified with these analyses (**Supplementary table 4**).

2) Post-surgery sample comparison between POD cases (n=10) vs non-POD controls (n = 27): There were no significant pathways identified with these analyses (**Supplementary table 5**).

<u>3) Pre-Post surgery sample comparison</u>

a) Pre-vs post-surgery sample comparison POD case and non-POD control together (n = 37)

Enrichment analysis showed pathways such as “response to lipopolysaccharide,” “inflammatory response,” and “interleukin-6 production” from GO analysis (**Supplementary table 6**), and “NF-kappa B signaling pathway” from KEGG analysis (**Supplementary table 6**), although those were not at the False Discovery Rate (FDR) significant level.

b) Pre-vs post-surgery sample comparison: POD case only (n = 10)

When only POD cases were tested for DNAm change before and after surgery, enrichment analysis showed many significant and intriguing pathways such as “cellular response to cytokine stimulus,” “response to cytokine,” “regulation of immune system process,” “regulation of cell activation,” and “regulation of cytokine production” from GO analysis (**Table 2**); and “HIF-1 signaling pathway,” “T cell receptor signaling pathway,” “Th17 cell differentiation,” “NF-kappa B signaling pathway,” and “Toll-like receptor signaling pathway” from KEGG analysis (**Table 2**), even at FDR significant level, other than the last few pathways from KEGG at the border of FDR significance (0.058) (**Table 2**).

**Table 2:**
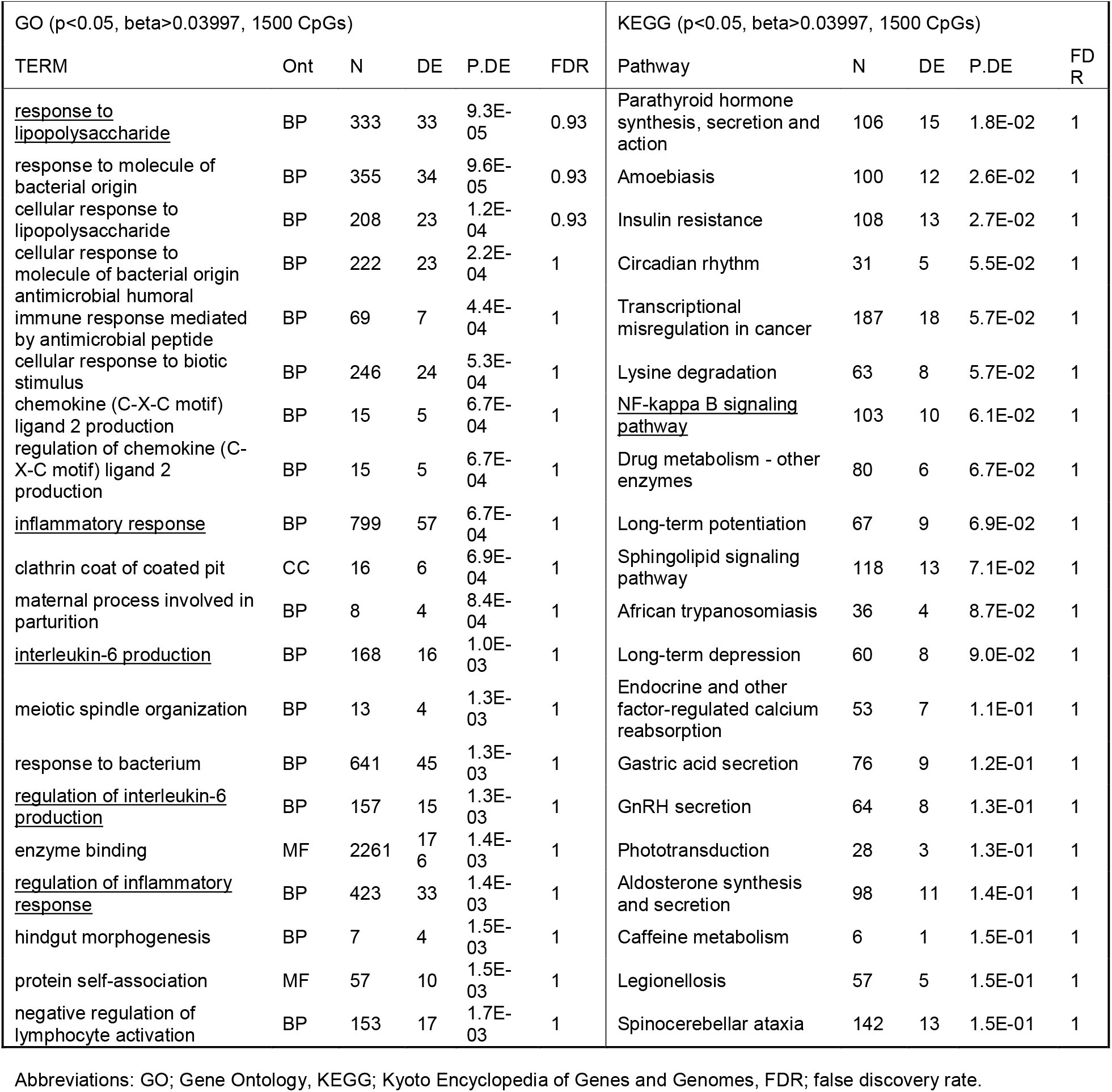
Result of the top 20 pathways of GO and KEGG analysis with differentially methylated CpGs between pre-surgery and post-surgery (all subjects)

**Table 2:** Result of the top 20 pathways of GO and KEGG analysis with differentially methylated CpGs between pre-surgery and post-surgery (POD group)

| GO (p < 0.05, beta > 0.03791, 1500 CpGs) |  |  |  |  |  | KEGG (p < 0.05, beta > 0.03791, 1500 CpGs) |  |  |  |  |
| --- | --- | --- | --- | --- | --- | --- | --- | --- | --- | --- |
| TERM | Ont | N | DE | P.DE | FDR | Pathway | N | DE | P.DE | FDR |
| <u>cellular response to cytokine stimulus</u> | BP | 1120 | 94 | 1.3E-08 | 2.9E-04 | <u>HIF-1 signaling pathway</u> | 109 | 21 | 3.2E-06 | 0.001 |
| <u>response to cytokine</u> | BP | 1210 | 98 | 3.2E-08 | 3.6E-04 | Kaposi sarcoma-associated herpesvirus infection | 193 | 26 | 4.1E-05 | 0.007 |
| cytokine-mediated signaling pathway | BP | 794 | 69 | 5.1E-08 | 3.9E-04 | EGFR tyrosine kinase inhibitor resistance | 78 | 17 | 1.3E-04 | 0.015 |
| <u>regulation of immune system process</u> | BP | 1512 | 116 | 8.9E-08 | 5.1E-04 | Inflammatory bowel disease | 63 | 11 | 2.7E-04 | 0.023 |
| <u>regulation of cell activation</u> | BP | 573 | 58 | 1.7E-07 | 7.6E-04 | Ras signaling pathway | 231 | 30 | 3.6E-04 | 0.024 |
| positive regulation of immune response | BP | 651 | 60 | 6.9E-07 | 0.002 | Prostate cancer | 97 | 17 | 4.7E-04 | 0.027 |
| T cell activation | BP | 482 | 51 | 7.3E-07 | 0.002 | Hepatitis B | 162 | 20 | 6.8E-04 | 0.031 |
| alpha-beta T cell activation | BP | 145 | 24 | 9.1E-07 | 0.002 | Human immunodeficiency virus 1 infection | 211 | 24 | 7.2E-04 | 0.031 |
| regulation of T cell activation | BP | 331 | 39 | 9.8E-07 | 0.002 | PD-L1 expression and PD-1 checkpoint pathway in cancer | 89 | 14 | 8.8E-04 | 0.033 |
| positive regulation of immune system process | BP | 985 | 81 | 1.2E-06 | 0.003 | <u>T cell receptor signaling pathway</u> | 102 | 16 | 0.001 | 0.034 |
| regulation of leukocyte activation | BP | 528 | 52 | 1.7E-06 | 0.003 | Insulin resistance | 108 | 16 | 0.001 | 0.035 |
| CD4-positive, alpha-beta T cell activation | BP | 97 | 18 | 1.9E-06 | 0.004 | Renal cell carcinoma | 68 | 12 | 0.002 | 0.055 |
| regulation of lymphocyte activation | BP | 440 | 46 | 2.0E-06 | 0.004 | <u>Th17 cell differentiation</u> | 105 | 15 | 0.002 | 0.055 |
| alpha-beta T cell differentiation | BP | 106 | 20 | 2.9E-06 | 0.005 | Human T-cell leukemia virus 1 infection | 216 | 25 | 0.002 | 0.055 |
| CD4-positive, alpha-beta T cell differentiation | BP | 78 | 16 | 3.0E-06 | 0.005 | <u>NF-kappa B signaling pathway</u> | 103 | 13 | 0.003 | 0.055 |
| <u>regulation of cytokine production</u> | BP | 790 | 62 | 4.0E-06 | 0.006 | Hepatocellular carcinoma | 168 | 21 | 0.003 | 0.055 |
| regulation of leukocyte cell-cell adhesion | BP | 328 | 36 | 5.9E-06 | 0.008 | Shigellosis | 243 | 25 | 0.003 | 0.055 |
| lymphocyte activation | BP | 687 | 62 | 6.2E-06 | 0.008 | B cell receptor signaling pathway | 80 | 12 | 0.003 | 0.058 |
| leukocyte cell-cell adhesion | BP | 363 | 38 | 9.2E-06 | 0.011 | Viral carcinogenesis | 193 | 21 | 0.003 | 0.058 |
| positive regulation of cytokine production | BP | 443 | 42 | 9.4E-06 | 0.011 | <u>Toll-like receptor signaling pathway</u> | 104 | 11 | 0.003 | 0.058 |

Abbreviations: GO; Gene Ontology, KEGG; Kyoto Encyclopedia of Genes and Genomes, FDR; false discovery rate.
Notes: The cut off for beta was determined to capture the top 1,500 CpG sites when p-value cut off was set at p<0.05. The underlines highlight the relevant pathways.

c) Pre-vs post-surgery sample comparison: non-POD control only (n = 27)

Enrichment analysis showed pathways such as “cell activation involved in immune response,” “neutrophil degranulation,” and “neutrophil-mediated immunity” from GO analysis, and “Longevity regulating pathway” from KEGG analysis, although those were not at the FDR significant level (**Table 3**).

**Table 3:**
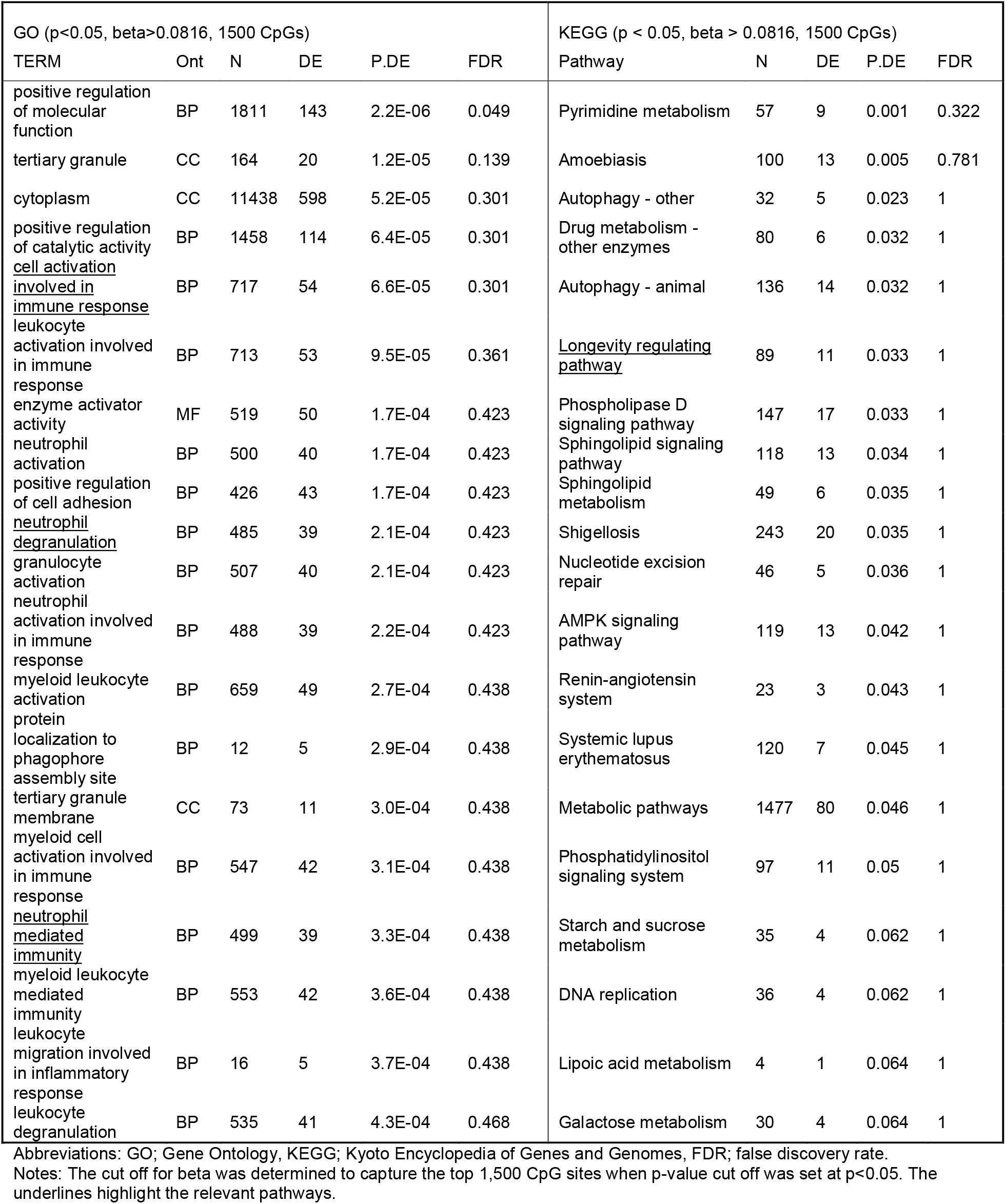
Result of the top 20 pathways of GO and KEGG analysis with differentially methylated CpGs between pre-surgery and post-surgery (non-POD group)

d) Pre-vs post-surgery sample comparison: POD case top CpG sites excluding overlapping non-POD control top CpG sites

To exclude the potential effect of common factors related to surgery and anesthesia between POD cases and non-POD controls, we excluded signals from non-POD controls to capture CpG sites associated to development of delirium after surgery. The top GO pathways from the enrichment analysis include “Immune response”, “immune system process”, “T cell activation”, and “regulation of immune system process” with FDR significant level. KEGG pathways showed “HIF-1 signaling pathway” and “T cell receptor signaling pathway” with FDR significant level (**Table 4**).

**Table 4:** Result of the top 20 pathways of GO and KEGG analysis with differentially methylated CpGs between pre- and post-surgery samples: POD case top CpG sites excluding overlapping non-POD control top CpG sites.

| GO (p<0.05, beta>0.03655, 1500 CpGs) |  |  |  |  |  | KEGG (p<0.05, beta>0.03655, 1500 CpGs) |  |  |  |  |
| --- | --- | --- | --- | --- | --- | --- | --- | --- | --- | --- |
| TERM | Ont | N | DE | P.DE | FDR | Description | N | DE | P.DE | FDR |
| <u>immune response</u> | BP | 1577 | 108 | 1.3E-06 | 0.030 | <u>HIF-1 signaling pathway</u> | 109 | 19 | 3.5E-05 | 0.012 |
| <u>immune system process</u> | BP | 2455 | 162 | 6.1E-06 | 0.046 | EGFR tyrosine kinase inhibitor resistance | 78 | 17 | 1.1E-04 | 0.014 |
| <u>T cell activation</u> | BP | 503 | 49 | 7.8E-06 | 0.046 | Prostate cancer | 97 | 18 | 1.3E-04 | 0.014 |
| <u>regulation of immune system process</u> | BP | 1358 | 100 | 8.0E-06 | 0.046 | Ras signaling pathway | 234 | 31 | 1.6E-04 | 0.014 |
| <u>leukocyte activation</u> | BP | 876 | 72 | 1.3E-05 | 0.060 | Kaposi sarcoma-associated herpesvirus infection | 194 | 24 | 2.4E-04 | 0.014 |
| <u>lymphocyte activation</u> | BP | 717 | 62 | 2.1E-05 | 0.081 | Inflammatory bowel disease | 63 | 11 | 2.5E-04 | 0.014 |
| <u>CD4-positive, alpha-beta T cell differentiation</u> | BP | 82 | 15 | 3.2E-05 | 0.102 | Hepatitis B | 162 | 20 | 6.1E-04 | 0.030 |
| <u>regulation of cell activation</u> | BP | 602 | 53 | 3.8E-05 | 0.102 | <u>T cell receptor signaling pathway</u> | 102 | 16 | 9.1E-04 | 0.038 |
| <u>alpha-beta T cell activation</u> | BP | 156 | 22 | 4.0E-05 | 0.102 | Hepatocellular carcinoma | 168 | 22 | 9.8E-04 | 0.038 |
| <u>cell activation</u> | BP | 1017 | 79 | 4.9E-05 | 0.111 | PD-L1 expression and PD-1 checkpoint pathway in cancer | 89 | 13 | 2.4E-03 | 0.084 |
| <u>alpha-beta T cell differentiation</u> | BP | 111 | 18 | 5.8E-05 | 0.113 | Viral carcinogenesis | 193 | 21 | 3.0E-03 | 0.088 |
| <u>leukocyte activation involved in immune response</u> | BP | 283 | 29 | 6.0E-05 | 0.113 | Human immunodeficiency virus 1 infection | 211 | 22 | 3.2E-03 | 0.088 |
| <u>CD4-positive, alpha-beta T cell activation</u> | BP | 101 | 16 | 6.4E-05 | 0.113 | <u>Toll-like receptor signaling pathway</u> | 104 | 11 | 3.3E-03 | 0.088 |
| <u>regulation of immune response</u> | BP | 810 | 62 | 7.5E-05 | 0.123 | AGE-RAGE signaling pathway in diabetic complications | 100 | 14 | 3.8E-03 | 0.088 |
| <u>intracellular signal transduction</u> | BP | 2640 | 192 | 8.7E-05 | 0.133 | Cellular senescence | 155 | 19 | 4.0E-03 | 0.088 |
| <u>cell activation involved in immune response</u> | BP | 287 | 29 | 9.5E-05 | 0.136 | African trypanosomiasis | 36 | 6 | 4.0E-03 | 0.088 |
| <u>regulation of leukocyte activation</u> | BP | 550 | 47 | 1.6E-04 | 0.221 | Acute myeloid leukemia | 67 | 11 | 4.9E-03 | 0.089 |
| <u>regulation of lymphocyte activation</u> | BP | 455 | 41 | 1.9E-04 | 0.237 | <u>Th17 cell differentiation</u> | 106 | 14 | 5.0E-03 | 0.089 |
| <u>adaptive immune response</u> | BP | 438 | 37 | 2.0E-04 | 0.237 | Shigellosis | 244 | 24 | 5.2E-03 | 0.089 |
| <u>cytidine to uridine editing</u> | BP | 12 | 4 | 2.2E-04 | 0.238 | Human cytomegalovirus infection | 225 | 24 | 5.4E-03 | 0.089 |

Abbreviations: GO; Gene Ontology, KEGG; Kyoto Encyclopedia of Genes and Genomes, FDR; false discovery rate.
Notes: The cut off for beta was determined to capture the top 1,500 CpG sites when p-value cut off was set at p<0.05. The underlines highlight the relevant pathways.

## Discussion

We conducted an epigenetics investigation of POD using genome-wide DNA methylation analysis. This is the very first study of its kind to the best of our knowledge. Our data identified significant evidence of numerous networks associated with POD, including response to cytokine, regulation of immune system process, cell activation and cytokine production, and positive regulation of immune response based on genome-wide DNAm signals from POD case blood comparing pre-versus post-surgery samples. Noteworthy is that although pre- and post-surgery comparison from combined total samples from both POD and non-POD cases showed certain signals associated with immune and inflammatory systems, such as response to lipopolysaccharide, inflammatory response, and IL-6 production, none of them met the FDR significance level. When limited to non-POD cases, pre- and post-surgery comparison did not show many relevant network signals even without the FDR significant level. Moreover, when signals from non-POD group were removed from POD group to exclude potential influence of surgery and anesthesia and to capture signals related to development of POD, the top signals showed immune response, T cell activation, T cell receptor signaling pathway, and HIF-1 signaling pathway. In fact, these top signals are very consistent with the top signals obtained from comparison of delirium patients versus non-delirium controls from independent inpatient cohort (15). This intriguing evidence of enriched signals among POD cases comparing post-surgery samples against pre-surgery samples suggests a dynamic epigenetics process not only induced through the surgery process but also influencing development of POD.

It has been suggested that systemic and brain inflammation play a key role in the development of delirium (28–31). Elevated levels of inflammatory markers in the blood and cerebrospinal fluid have been demonstrated in patients with POD (32). It is well known that peripherally produced cytokines act on the brain to cause abnormal behaviors because the immune-to-brain communication pathways leads to the pro-inflammatory cytokines production by microglial cells (33). Indeed, animal studies have demonstrated that peripheral lipopolysaccharide injection or surgical intervention, such as laparotomy or hepatectomy, to rodents induced cognitive deficit–related behavior along with heightened inflammatory cytokines such as IL-1beta, IL-6, TNF-alpha in the brain (34). With regard to epigenetics influence from surgery, a recent study by Sadahiro et al. also demonstrated that various major surgeries cause changes in DNAm at sites annotated to immune system genes, which is consistent with our present data (35). The report did not have information about the differences in DNAm between POD and non-POD (35). Thus, our study further advanced the understanding of pathophysiology of delirium from an epigenetic stand point. It can be suggested that alterations in DNAm of genes related to inflammation and immunity in POD patients observed in our current enrichment analyses are involved in the peripheral inflammation and immunity, as well as subsequent neuroinflammation and onset of delirium.

There are several strengths to this study. First, the surgery type was only neurosurgery for medication-refractory epilepsy. Thus, our study participants experienced relatively similar insult across the cohort. This is a definite advantage over our previous approach using inpatients from diverse background and etiologies of delirium (15–17). Second, it is important to mention that the pre-post surgical sample collection design allowed us to conduct pre-post comparison from same individuals to directly measure the impact of surgery on their DNAm signature and to find differences between POD cases vs non-POD controls. The post surgical blood was collected immediately after the surgery, which was the time before the onset of POD. Therefore, we were able to see differences in epigenetics status related to POD based on blood data even before the onset. We believe that such information based on blood identified after surgery but before onset of POD is useful for future clinical practice, as such biomarker can potentially give us an opportunity to identify high risk patients even before the emergence of symptoms.

Such data can help us further identify a potential epigenetics mechanism in how exogenous insults influence the epigenetic landscape leading to development of delirium. Through this design, it is feasible to separate out DNAm change induced by the surgical process including anesthesia, and DNAm change associated with—and potentially responsible for—development of delirium. To support the notion, the top pathways identified with our analysis was exactly consistent with the top pathways we previously reported from independent non-surgical inpatient cohort. Third, DNAm marks from pre-surgery samples can be used as a potential predictive biomarker for POD. Although our present data did not show significant difference from pre-surgery samples, it is still possible that with larger sample size we may find signals usable for prediction of POD. If such data is replicated and confirmed, an algorithm based on a pre-surgical DNAm signature can help identify patients at high risk for POD, and appropriate intervention and prevention can be employed. Overall, the data presented here are consistent with our hypothesis and support an epigenetics mechanism in the pathophysiological process of delirium.

We acknowledge several limitations in this study as follows. First, the sample size is small, limited to 37 subjects total and only 10 POD patients in this cohort. However, availability of pre-post surgery blood samples from same individuals are a merit of this investigation. Second, it is possible that other surgery types or other insults such as infection may induce a different epigenetics process from the pathways we identified in our specific cohort of neurosurgery patients. However, our previous data from a more diverse group of inpatients with and without delirium from various etiologies showed overlapping signals, suggesting that our present findings have at least some levels of generalizability to other populations with delirium. Third, in our cohort, age range is not limited to elderly subjects. Also, average age between POD and non-POD groups were different. To address the potential influence of age difference, we included age as a covariate in our analysis to minimize the influence. Once the sample size is increased in future study, it is expected that we will be able to investigate the potential age-dependent effect on our findings reported here. Fourth, there was no single gene or CpG site identified as significant at the genome-wide level, although it was expected given the small sample size. Thus, selection of CpG sites for downstream enrichment analysis requires a careful approach. Nonetheless, the enrichment analysis from genome-wide DNAm data showed strong evidence of relevant pathways playing a role in POD. Fifth, our data merely shows the association between epigenetics signals and POD and does not suggest a causal relationship. However, blood samples were collected just after surgical procedure in the OR, before the actual development of POD. Thus, it is reasonable to believe that such DNAm changes occur before the development of POD, indicating that certain epigenetics process may be playing a role in the pathophysiological mechanism of POD, instead of POD causing such DNAm changes. Sixth, we conducted this study at a single institution, and most of study subjects were non-Hispanic white. Therefore, generalizability needs to be confirmed with more diverse ethnic groups. Seventh, our definition of POD relied solely on a retrospective chart review of electronic medical records. Therefore, it is possible that there are false positive or false negative cases in our data set. However, even with the possibility of misclassification, we observed several intriguing and relevant signals as described above.

In summary, this is the first study demonstrating the differences of DNAm profiles between POD and non-POD patients using genome-wide methylation analysis. Our findings provide further evidence of the potential role of epigenetics in the pathophysiology of delirium.

## Funding

This work was supported by research grants from the National Institute of Mental Health, United States (R01 MH119165). Gen Shinozaki receives research grant support from National institute of health (NIH), National Science Foundation (NSF), Sumitomo Pharma LTD, and Fujitsu Laboratories LTD.

## Role of the Sponsors

The supporters had no role in the design, analysis, interpretation, or publication of this study.

## Declaration of Interest

Gen Shinozaki has pending patents “Epigenetic Biomarker of Delirium Risk” in the PCT Application No. PCT/US19/51276, in the PCT Application No. PCT/US21/63166, and in U.S. Provisional Patent No. 62/731,599. All other authors have declared that no conflict of interest exists.

## Acknowledgment

The authors thank the patients who participated in this study. Also, we would like to thank Paul J. Casella for English language editing. This work was supported by research grants from the National Institute of Mental Health, United States (R01 MH119165).

## Author information

Takehiko Yamanashi, Kaitlyn J. Crutchley, and Nadia E. Wahba contributed equally. These three co-first authors can prioritise their names when adding paper’s reference to their resumes.

## Author Contributions

T.Y., K.J.C., and N.E.W. analyzed data and wrote the initial draft of the manuscript. P.S.M. and H.R.C. analyzed data. T.N. collected samples and clinical data. C.C.A. and E.J.S. processed samples. H. K. participated in its design, collected samples, and helped coordination. M.I., M.A.H., C.G.H., P.P., and M.M.H critically reviewed the manuscript. G.S. conceived ideas of the study, planned its design and coordination, drafted the initial form of the manuscript and edited the manuscript for the final form.

## Data Availability

The data that support the findings of this study are available from the corresponding author, G.S., upon reasonable request.

