## Supplementary figures and images for "Epigenetics of post-operative delirium: A genome-wide DNA methylation study of neurosurgery patients"

### Supplementary Figure

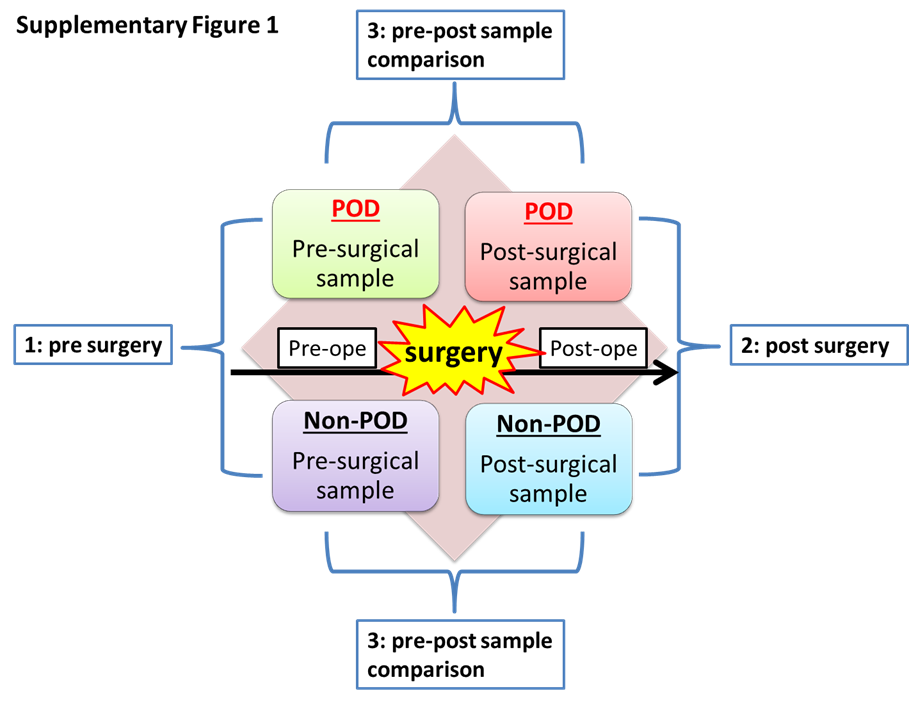
